# P2Y_6_ signaling selectively regulates TLR3-induced chemokine release in murine bone marrow-derived macrophages, with opposing effects on CCL2 and CXCL1

**DOI:** 10.64898/2026.09.13.751193

**Authors:** Fatemeh Salarpour, Anaïs Brafine, Ashley Charre, Julie Pelletier, Jean Sévigny

## Abstract

Extracellular nucleotides regulate diverse inflammatory responses through the activation of P2 receptors. Here, we show that nucleotides regulate TLR3-induced chemokine responses in murine bone marrow-derived macrophages (BMDMs). Stimulation with the TLR3 agonist, poly(I:C), induced CCL2 and CXCL1 secretion, whereas degradation of extracellular nucleotides with apyrase reduced CCL2 but enhanced CXCL1 release. In agreement, the non-selective P2 receptor antagonists suramin and RB2 reduced poly(I:C)-induced CCL2 production. Among the highly expressed P2 receptors examined, selective inhibition of P2Y_6_ with MRS2578 reproduced the effects of apyrase, decreasing CCL2 while increasing CXCL1 secretion. The absence of effects of MRS2578 in *P2ry6*-deficient BMDMs confirmed the specificity of this antagonist. Transcriptional analysis further revealed that P2Y_6_ inhibition reduced early poly(I:C)-induced *Ccl2* and *Cxcl10* expression while enhancing later *Cxcl1* and *Cxcl2* expression. In line with our previous works on other pattern-recognition receptors pathways and innate immune cells, this study further supports a role for extracellular nucleotides in shaping TLR-induced chemokine responses in macrophages.

## Introduction

Macrophages play a key role in innate immune defense against invading pathogens through the activation of pattern-recognition receptors (PRRs), including Toll-like receptors (TLRs). Macrophages express various TLRs including TLR2 (activated by peptidoglycan and lipopeptide), TLR3 (activated by double-stranded RNA within endosomal compartments), TLR4 (activated by LPS), TLR5 (activated by the bacterial flagellin) and TLR7/8 (activated by single-stranded viral RNA)(1–4). In response to TLR activation, macrophages release cytokines and chemokines that coordinate innate immune responses. Two important examples are C-C motif chemokine ligand 2 (CCL2; MCP-1), which acts primarily through C-C chemokine receptor 2 (CCR2) to recruit inflammatory monocytes, and C-X-C motif chemokine ligand 1 (CXCL1; KC), which signals through C-X-C chemokine receptor 2 (CXCR2) to promote neutrophil recruitment (5,6).

In addition to TLRs, macrophages express P2 receptors that are activated by extracellular nucleotides (7). P2 receptors are divided into two families: G protein-coupled P2Y receptors and ligand-gated P2X ion channels. These receptors differ in their nucleotide selectivity, allowing extracellular ATP, ADP, UTP, and UDP to activate distinct signaling pathways (8). Previous studies have reported the high expression of several P2 receptors, including P2X4,7 and P2Y_1,2,6,14_ subtypes in macrophages (7). Other studies have reported that P2Y_6_ regulates cytokine and chemokine production in mouse macrophages (9), P2X4 promotes PGE2 release from tissue-resident macrophages (9), and P2X7 mediates IL-1β release and macrophage cell death (10).

There is growing evidence that extracellular nucleotides regulate several immune functions, including cytokine and chemokine production, migration, phagocytosis, inflammasome activation, and cell survival (11,12). They also appear to contribute to inflammatory responses initiated by PRRs. Several studies have demonstrated that extracellular nucleotides participate in TLR4-induced inflammatory responses. In addition, we previously demonstrated that activation of TLR1/2 with Pam3CSK4 in human primary monocytes and monocytic cells induce CXCL8/IL-8 production through autocrine stimulation of P2Y_2_ and P2Y_6_ receptors (13). We also showed that extracellular nucleotide degradation with apyrase and P2Y_6_ inhibition reduced LPS/TLR4-induced CXCL8 and CCL2 secretion in human glioma cells (14). In mouse macrophages, UDP–P2Y_6_ signaling was shown to potentiate LPS/TLR4-induced IL-6 and CXCL2/MIP-2 production, further supporting a role for extracellular nucleotides in TLR-induced inflammatory responses (15).

In this work, we show that extracellular nucleotides also regulate chemokine production induced by the TLR3 agonist Polyinosinic-polycytidylic acid (poly I:C) in murine BMDMs.

## Materials & Methods

Poly(I:C) was purchased from InvivoGen (San Diego, California, USA). Potato apyrase, suramin and uridine 5’-diphosphate (UDP) were purchased from Sigma–Aldrich (Oakville, ON, Canada or St. Louis, MO, USA). MRS2578 (P2Y_6_ antagonist), 5-BDBD (P2X4) and A-438079 (P2X7) were purchased from Tocris Bioscience (Bristol, UK), and DMSO from Sigma Chemical. M-CSF were purchased from PeproTech (QC, Canada). FBS (heat-inactivated by a 30 min incubation at 56°C), DMEM/F12, L-glutamine, penicillin and streptomycin, and Dulbecco’s phosphate-buffered saline (D-PBS) containing calcium and magnesium (Cat. No. 311-430-CL) were purchased from Wisent Inc. (Saint-Jean-Baptiste, QC, Canada). Mouse CXCL1/KC and CCL2/MCP-1 were purchased from R&D Systems (Minneapolis, MN, USA). Fluorochrome-conjugated monoclonal antibodies against mouse were purchased from BioLegend (San Diego, CA, USA). All reagents were reconstituted according to the manufacturers’ instructions and stored at −20°C or −80°C as recommended.

### Animals

All animal protocols were approved by the Université Laval Animal Care Committees and followed the Canadian Council on Animal Care (CCAC) guidelines. Wild-type (WT) C57BL/6 and *P2ry6*^−*/*−^ mice were derived from a specific-pathogen-free elite colony and bred in the animal facility of the Centre de recherche du CHU de Québec–Université Laval. The *P2ry6*^−*/*−^ mice had been backcrossed onto the C57BL/6 background for 14 generations (15). Adult male WT C57BL/6 mice (8–12 weeks old), either bred in-house or purchased from Charles River Laboratories (Pointe-Claire, QC, Canada), were used as controls. Animals were housed under specific-pathogen-free conditions in a temperature-controlled room (21°C) with a 12 h light/12 h dark cycle and had unrestricted access to a standard diet and tap water, unless otherwise specified.

### Generation of bone marrow-derived macrophages (BMDMs)

Bone marrow cells were isolated from the femurs and tibias of WT or *P2ry6*-deficient mice. Briefly, the epiphyses were removed, and the bones were placed in a perforated 0.5-mL microcentrifuge tube nested inside a 1.5-mL microcentrifuge tube and centrifuged at 10,000 × g for 30 s to expel the bone marrow. The collected bone marrow cells were resuspended in sterile D-PBS, supplemented with 1% (v/v) FBS, passed through a 70-µm cell strainer to obtain a single-cell suspension, and centrifuged at 500 × g for 10 min. The cell pellet was resuspended in complete macrophage medium consisting of DMEM/F12 supplemented with 10% (v/v) FBS, 100 U/mL penicillin, 100 µg/mL streptomycin, 10mM L-glutamine, and 10 ng/mL recombinant macrophage colony-stimulating factor (M-CSF). Cells were seeded at appropriate density directly in tissue culture plates and cultured in complete macrophage medium at 37°C in a humidified incubator with 5% CO_2_ for 7 days to allow differentiation into BMDMs. Fresh medium containing M-CSF was added on day 3. On day 7, cells were washed two times with D-PBS to remove non-adherent cells, and the adherent BMDMs were used directly for subsequent experiments (16,17).

### BMDMs stimulation and conditioned media

For stimulation experiments, BMDMs were generated directly in 24-well tissue culture plates at a density of 0.5 × 10^6^ cells/well in complete macrophage medium supplemented with 10 ng/mL M-CSF. On day 7, the differentiated adherent BMDMs were stimulated with poly(I:C) (0.1 or 10 µg/mL) in the presence or absence of compounds added 20 min before stimulation: the nucleotide scavenger apyrase (2 U/mL); the non-selective P2 receptor antagonists suramin (100 µM) and Reactive Blue 2 (RB2; 100 µM); or the selective P2 receptor antagonists MRS2578 (P2Y_6_, 5 µM), 5-BDBD (P2X4, 5 µM), and A-438079 (P2X7, 5 µM). Vehicle-treated cells received 0.01% DMSO, corresponding to the final concentration used for the selective antagonists. Where indicated, uridine 5′-diphosphate (UDP; 100 µM) was added simultaneously aspoly(I:C). For chemokine release and gene expression analyses, cells were stimulated for 18 h and for 1 or 5 h, respectively. Cell viability after treatments was monitored in representative experiments by trypan blue exclusion to ensure that changes in chemokine production reflected regulated responses rather than cytotoxicity.

### Enzyme-linked immunosorbent assay (ELISA)

Following stimulation, cell culture supernatants were collected and stored at −80°C until analysis. Concentrations of CXCL1 and CCL2 were quantified using DuoSet ELISA kits (R&D Systems Inc., Minneapolis, MN, USA), according to the manufacturer’s instructions. Briefly, high-binding 96-well plates were coated overnight at 4°C with capture antibody, blocked, and incubated with appropriately diluted samples and CXCL1 or CCL2 standards. Standard curves were generated using recombinant mouse CXCL1 or CCL2 to enable absolute quantification. After incubation with detection antibody and streptavidin-HRP, plates were developed with TMB substrate, and the reaction was stopped with 2N H_2_SO_4_. Absorbance was measured at 450 nm with wavelength correction at 570 nm using a microplate reader. Chemokine concentrations were calculated from standard curves generated with recombinant mouse CXCL1 or CCL2.

### RNA extraction and quantitative RT-PCR

Total RNA was isolated from unstimulated and stimulated BMDMs using TRIzol reagent (Invitrogen, Carlsbad, CA, USA) according to the manufacturer’s instructions and quantified spectro-photometrically at 260 nm. Complementary DNA (cDNA) was synthesized from 1 µg of total RNA using SuperScript III reverse transcriptase and oligo (dT) primers, following the manufacturer’s instructions (Invitrogen, Carlsbad, CA, USA). Gene expression was assessed by qPCR using SYBR Green Supermix (Roche Diagnostics, Mannheim, Germany) and primers specific for *Gapdh, P2ry6, P2rx4, P2rx7* and *Tlr3* receptors and chemokines including *Cxcl1, Cxcl2, Ccl2* and *CXCL10*. Primers were either designed in-house and synthesized by Invitrogen (Carlsbad, CA, USA) or purchased from Qiagen (Toronto, ON, Canada), as listed in Table 1. Standard curves were used to determine mRNA transcript copy number in individual reactions. *Gapdh* was used to normalize RNA quantities between samples.

**Table 1.**
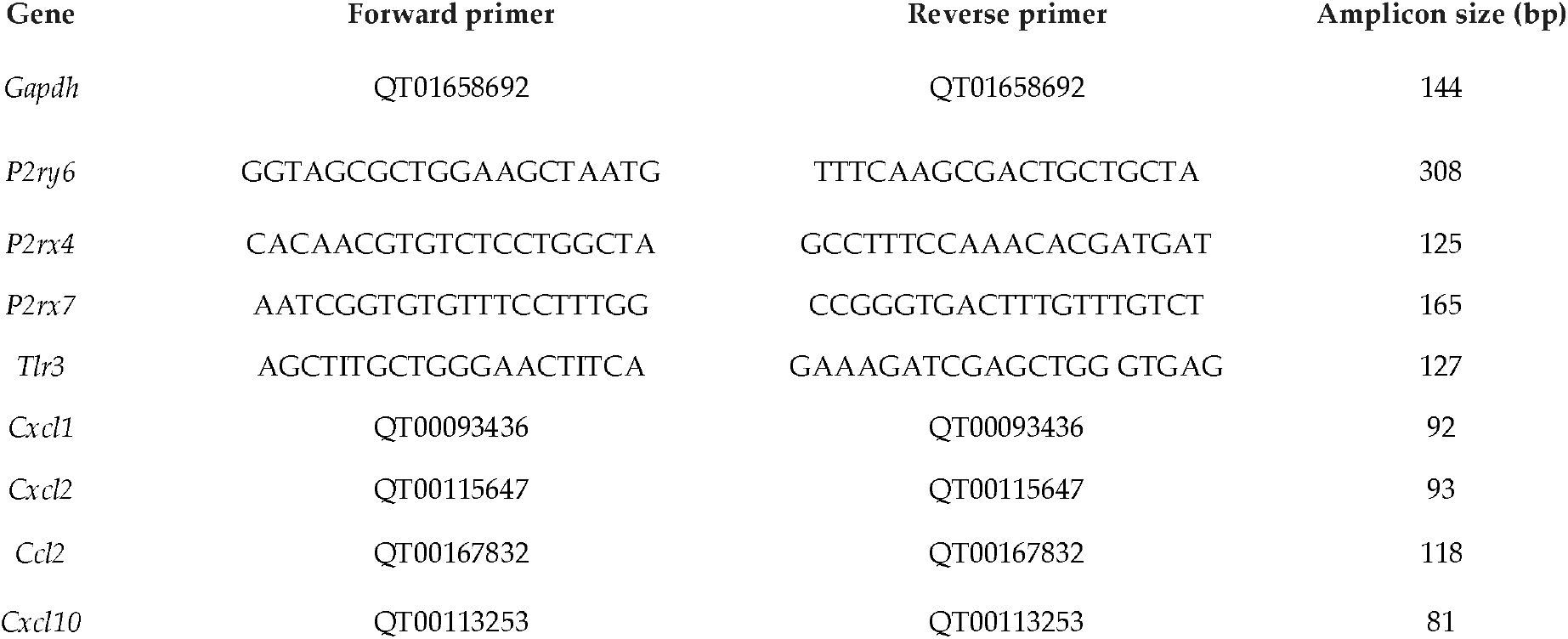
Primer pairs for PCR and amplicon size.

### Reanalysis of public RNA-seq data

Publicly available RNA-sequencing data were retrieved from the Gene Expression Omnibus (GEO) database (accession GSE198821), with raw sequencing reads obtained from the corresponding Sequence Read Archive (SRA) BioProject (PRJNA817065) (18). RNA-seq data were analyzed using the Galaxy platform. Raw FASTQ files were downloaded using the SRA Toolkit (fasterq-dump). Sequence quality was evaluated using FastQC and summarized with MultiQC. Reads were mapped to the Mus musculus reference genome (GRCm39) using STAR with the corresponding Ensembl genome sequence and gene annotation. Gene-level read counts were generated using HTSeq-count. Raw count data were loaded into R (version 4.6.1) and analyzed using the edgeR package. Library sizes were normalized using the trimmed mean of M-values (TMM) method, and normalized gene expression values were reported as counts per million (CPM). Bar plots were created using normalized CPM values. Genes involved in nucleotide signaling and inflammatory pathways were selected for downstream analyses.

### Flow cytometry

BMDMs were harvested (1 × 10^6^ cells/condition) and stained with fluorochrome-conjugated antibodies against CD11b (PerCP-Cy5.5) and F4/80 (PE) following Fc receptor blocking. Dead cells were excluded using AmCyan-A viability dye. Samples were acquired using a flow cytometer (BD FACS Canto™ II) and analyzed with Flow Jo software. Macrophages were identified after sequential gating on singlets, viable cells, and CD11b^+^F4/80^+^ populations. Cells were sequentially gated to exclude debris based on forward scatter (FSC) and side scatter (SSC) characteristics, followed by singlet discrimination and exclusion of non-viable cells. BMDMs were identified as F4/80^+^ and CD11b^+^.

### Statistical analysis

Data are presented as the mean ± SEM from at least three independent experiments unless otherwise indicated. Statistical analyses were performed using GraphPad Prism (10, 2025). Comparisons between two groups were performed using Student’s T-test. Multiple-group comparisons were analyzed by one-way ANOVA. A *p* value < 0.05 was considered statistically significant.

## Results

### Flow cytometric characterization of WT and P2ry6-deficient BMDMs

To confirm the cellular model, BMDMs generated from WT and *P2ry6*-deficient mice were characterized by flow cytometry. Cells were sequentially gated based on FSC/SSC characteristics to exclude debris, followed by singlet discrimination and exclusion of non-viable cells. Macrophages were identified as CD11b^+^F4/80^+^ cells. M-CSF differentiation yielded highly enriched macrophage cultures, with more than 95% of viable singlet cells exhibiting the CD11b^+^F4/80^+^ phenotype in both WT and *P2ry6*-deficient cultures (Fig. 1A & B). These results confirm efficient BMDM differentiation and show that the proportion of differentiated macrophages was comparable between WT and *P2ry6*-deficient BMDMs.

**Figure 1.**
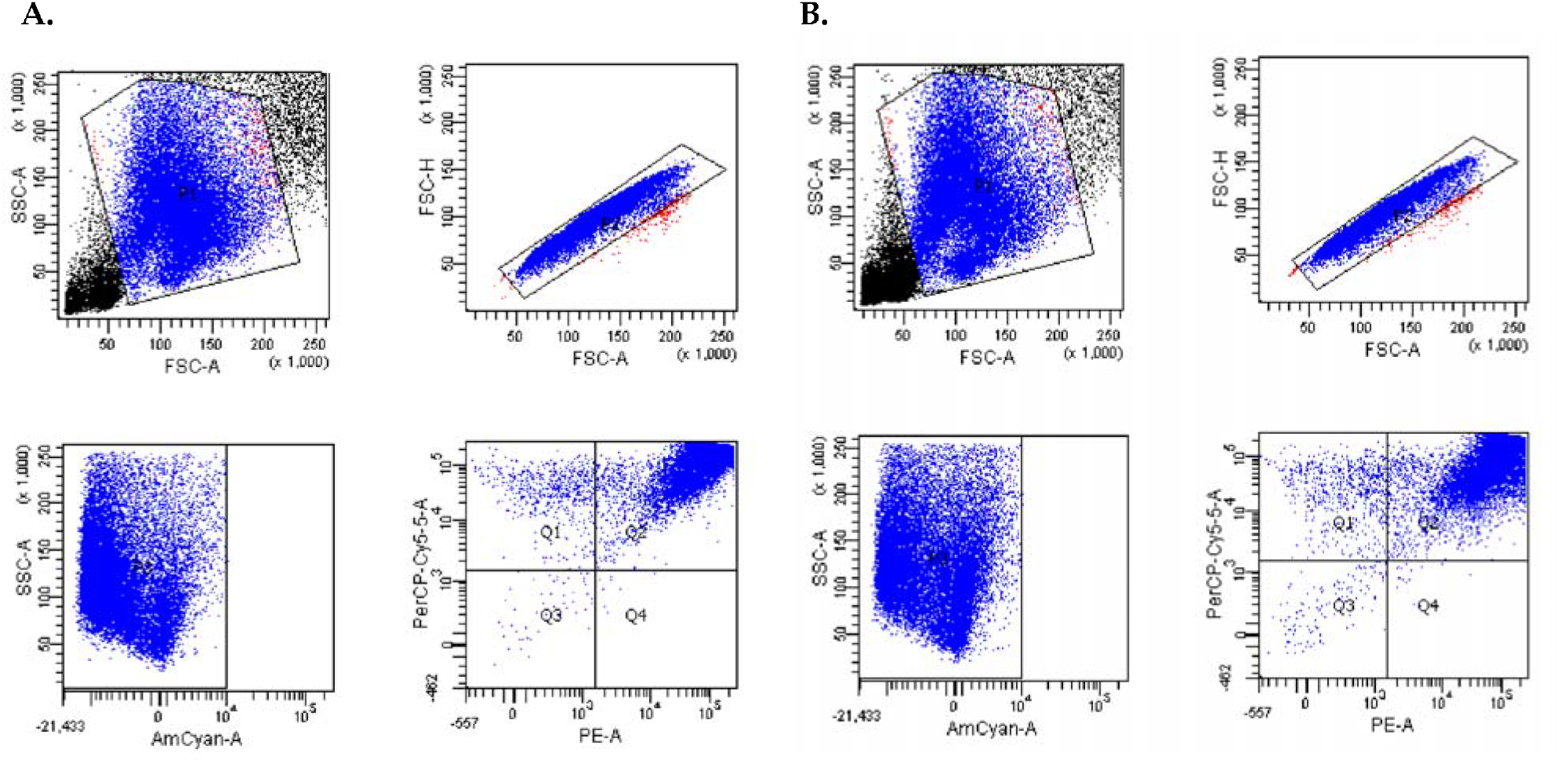
Flow cytometric characterization of WT and *P2ry6*-deficient murine BMDMs. Representative flow-cytometric analysis of BMDMs generated from WT (Panel A) and *P2ry6*-deficient (Panel B) mice. Cells were sequentially gated according to forward- and side-scatter characteristics (FSC-A/SSC-A) to exclude debris, followed by singlet discrimination using FSC-A/FSC-H and exclusion of non-viable cells using an AmCyan-A viability dye. Macrophages were identified as CD11b^+^F4/80^+^ cells, with CD11b detected using PerCP-Cy5.5 and F4/80 detected using PE. More than 95% of viable singlet cells displayed the CD11b^+^F4/80^+^ macrophage phenotype. All plots are from the same representative sample selected from three independent experiments performed with different mice.

### TLR3 mediates CCL2 and CXCL1 release in BMDMs

We first assessed the basal expression of *Tlr3, Ccl2*, and *Cxcl1* by reanalysis of publicly available RNA-seq data (Fig. 2A) (18), and subsequently confirmed their expression by RT-qPCR in our BMDMs (Fig. 2B). Given the important role of TLR3 in macrophage inflammatory responses (19) and of the involvement of (19) extracellular nucleotide signaling (17,20), BMDMs were next stimulated for 18 h with increasing concentrations of poly(I:C), after which CCL2 (Fig. 2C) and CXCL1 (Fig. 2D) levels in the culture supernatants were quantified by ELISA. These chemokines were selected as CCL2 and CXCL1 are major mediators of monocyte and neutrophil recruitment, respectively (21,22). poly(I:C) induced a concentration-dependent increase in CCL2 and CXCL1 secretion, reaching maximal effects at 10 µg/mL.

**Figure 2.**
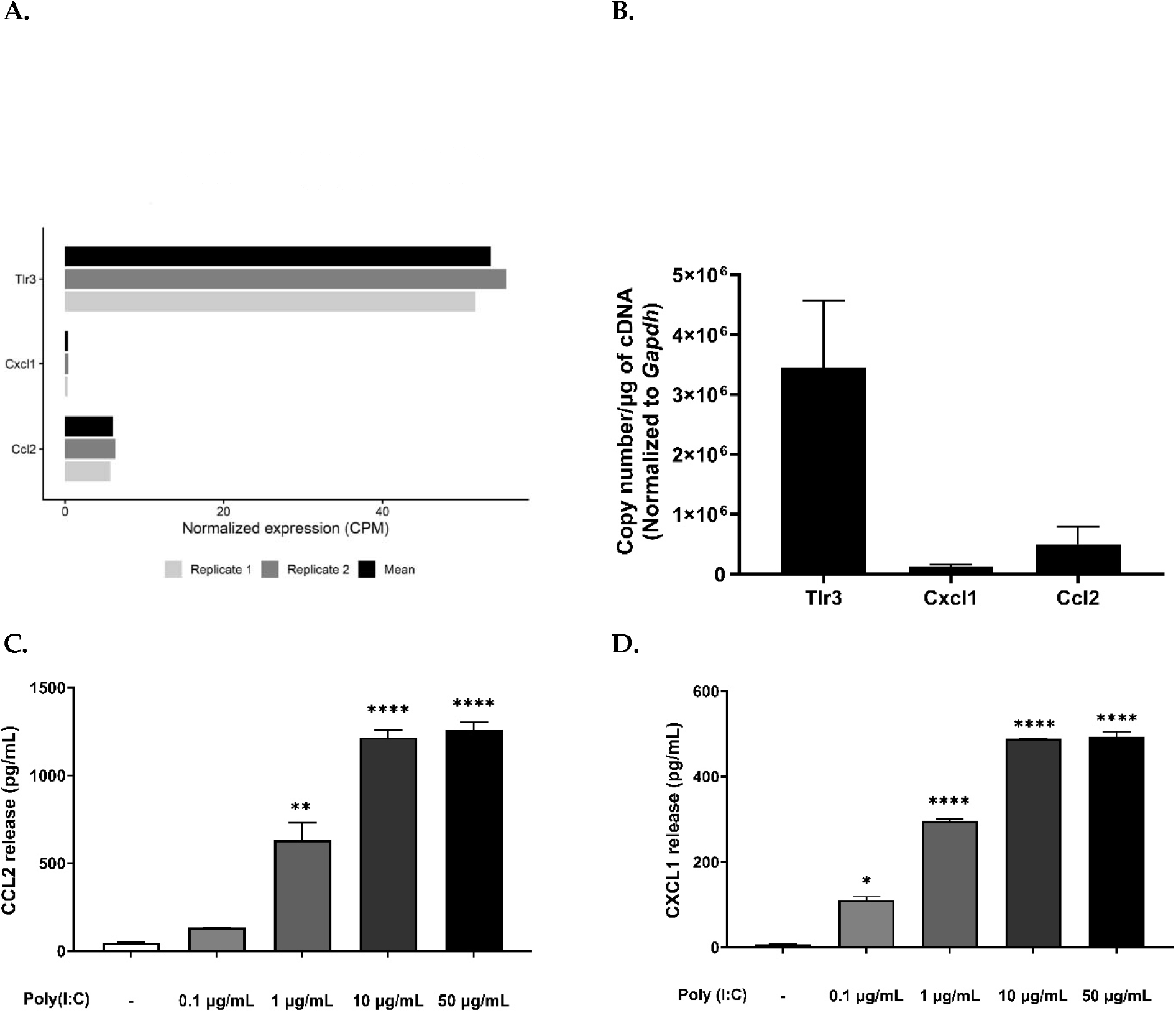
Poly(I:C) induces CCL2 and CXCL1 release in BMDMs. Basal expression of *Tlr3, Ccl2* and *Cxcl1* in unstimulated BMDMs (Panel A) by reanalysis of publicly available RNA-seq data from GSE198821 (PRJNA817065), expressed as counts per million (CPM). Basal expression of those genes was additionally assessed by RT-qPCR in unstimulated BMDMs and normalized to *Gapdh* (Panel B). Poly(I:C) stimulation induces CCL2 (Panel C) and CXCL1 (Panel D) release in a concentration-dependent manner in BMDMs. Cells were stimulated with poly(I:C) at 50, 10, 1 and 0.1 µg/mL for 18 h and chemokine levels in culture supernatants were quantified by ELISA. Data are presented as mean ± SEM of three independent experiments. Each condition was conducted in triplicate. One symbol indicates *p* < 0.05, two symbol p < 0.01 and four symbols p < 0.0001. *, poly(I:C)-stimulated cells compared with unstimulated cells.

### Extracellular nucleotides are involved in poly(I:C)-induced CCL2 and CXCL1 release in BMDMs

In previous work on myeloid cells, we observed that the activation of TLR2, TLR4 in human monocytes induced CXCL8 secretion during concomitant P2Y_2_ and P2Y_6_ receptor engagement (13)). We also observed reduced CXCL10 expression and secretion following P2Y_6_ inhibition in intestinal epithelial cells (17). These observations led us to investigate whether a similar nucleotide-dependent mechanism functions downstream of TLR3 in BMDMs. Supporting this hypothesis, poly(I:C)-stimulated BMDMs treated with the nucleotide scavenger apyrase or the broad P2 antagonists suramin and RB2 exhibited significantly suppressed CCL2 release (Fig. 3A) alongside enhanced CXCL1 secretion (Fig. 3B). Together, these findings link extracellular nucleotide signaling to poly(I:C)-induced chemokine secretion in BMDMs and demonstrate that extracellular nucleotides differentially regulate these chemokine responses.

**Figure 3.**
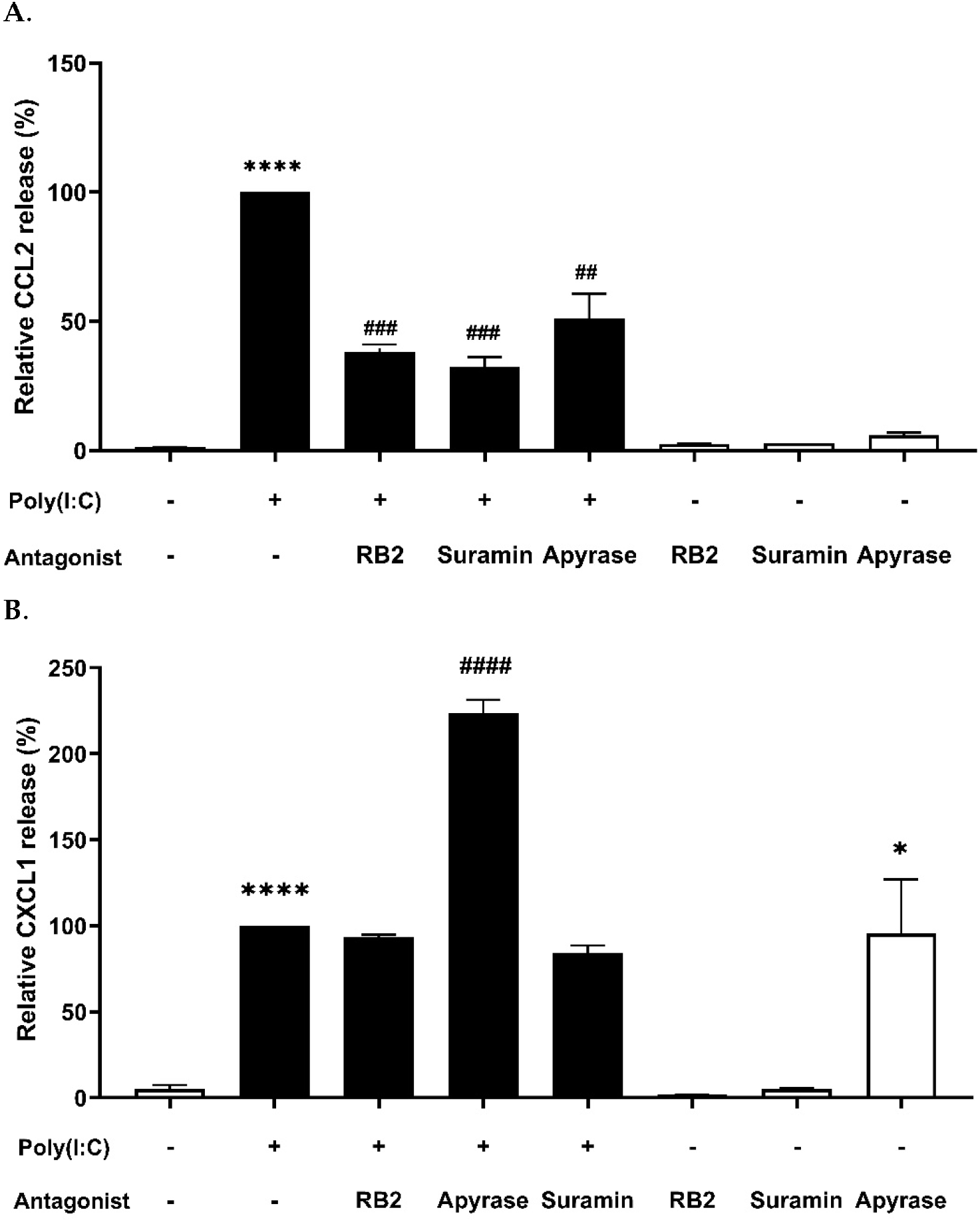
Extracellular nucleotides are involved in poly(I:C)-induced CCL2 and CXCL1 release in BMDMs. Cells were stimulated with poly(I:C) (10 µg/mL) for 18 h in the presence or absence of the nucleotide scavenger apyrase (2 U/mL), the non-selective P2 receptor antagonist suramin (100 µM), or Reactive Blue 2 (RB2) (100 µM), added 20 min before stimulation. The release of CCL2 (A) and CXCL1 (B) in the supernatants was quantified by ELISA. Data are presented as the mean ± SEM of three independent experiments, each performed with BMDMs from a different mouse. The 100% values correspond to absolute CCL2 and CXCL1 concentrations of 1250 and 500 pg/mL, respectively, following stimulation with poly(I:C) at 10 µg/mL (Panel A & B), as reported in Fig. 2 C & D. Two symbol p < 0.01, three symbols p < 0.001, and four symbols p < 0.0001. *, poly(I:C)-stimulated cells compared with unstimulated cells; #, poly(I:C)-stimulated cells in the presence of inhibitors compared with poly(I:C) alone.

### P2Y_6_ receptors are required for poly(I:C)-induced CCL2 and CXCL1 secretion in BMDMs

We next investigated which P2 receptors contributed to poly(I:C)-mediated CCL2 and CXCL1 release. Reanalysis of the RNA-seq dataset revealed a distinct P2 receptor expression profile in unstimulated BMDMs, with relatively high expression of *P2ry6, P2rx4*, and *P2rx7* compared with the other P2 receptors examined (Fig. 4A, B) (18). Expression of these three receptors was confirmed by RT-qPCR in BMDMs generated in our experimental system (Fig. 4C). Based primarily on their relatively high expression in BMDMs, *P2ry6, P2rx4*, and *P2rx7* were therefore selected as candidate receptors for functional investigation. This selection was also consistent with previous studies demonstrating functional roles for these receptors in macrophage inflammatory responses (9,15,23). BMDMs were then stimulated with Poly(I:C) (10 µg/mL) in the presence or absence of selective antagonists targeting P2Y_6_ (MRS2578), P2X4 (5-BDBD), or P2X7 (A-438079). Among the receptors tested, only P2Y_6_ inhibition significantly altered the Poly(I:C)-induced chemokine response. This receptor inhibition reduced CCL2 release by approximately 40% (Fig. 4D), whereas it increased CXCL1 secretion by about 50% under the same conditions (Fig. 4E). However, the inhibition of P2X4 or P2X7 did not significantly affect the secretion of either chemokine. These findings identify P2Y_6_ as the main nucleotide receptor regulating poly(I:C)-induced chemokine release in BMDMs under the experimental conditions tested and reveal that its contribution differs between CCL2 and CXCL1.

**Figure 4.**
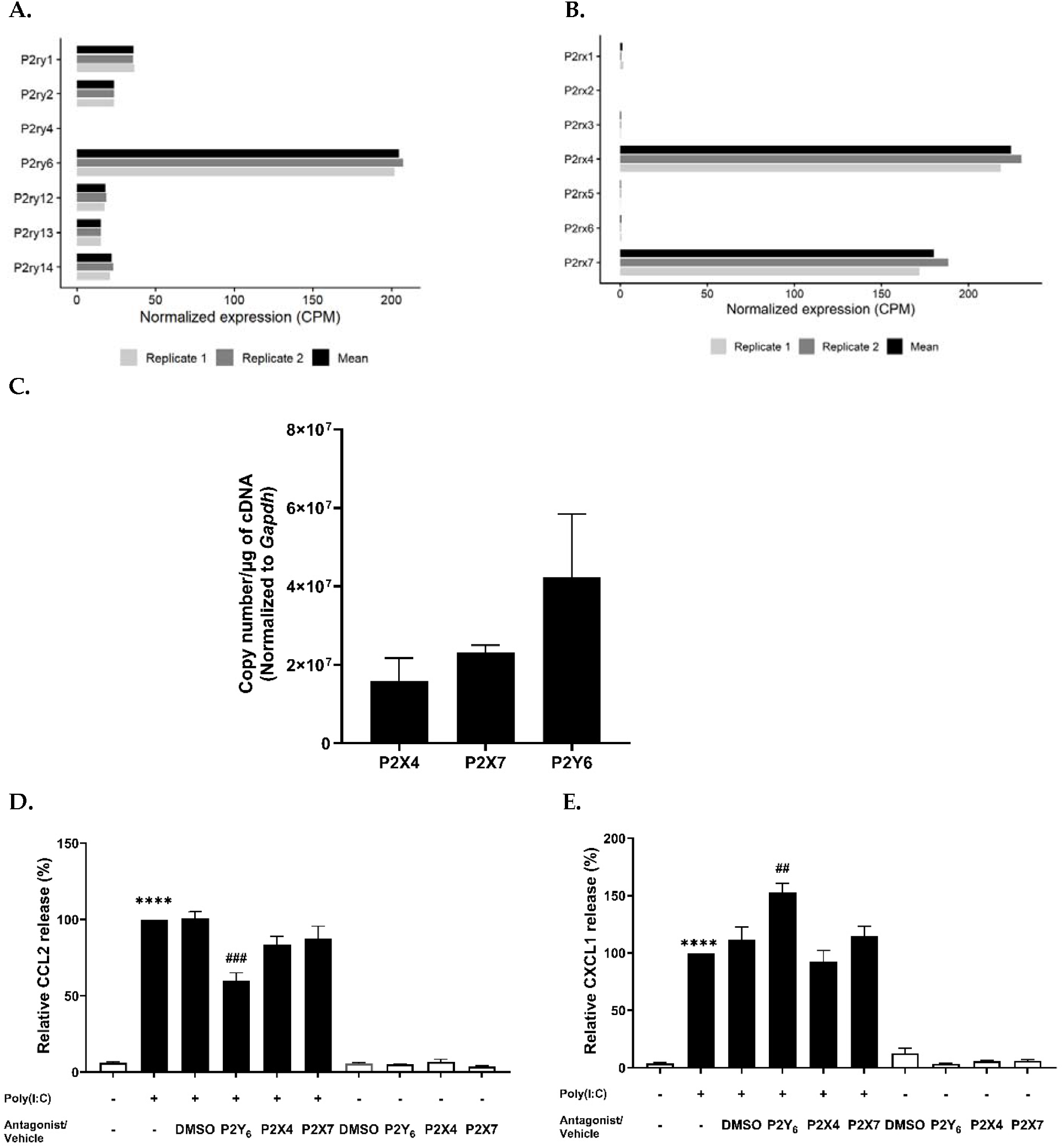
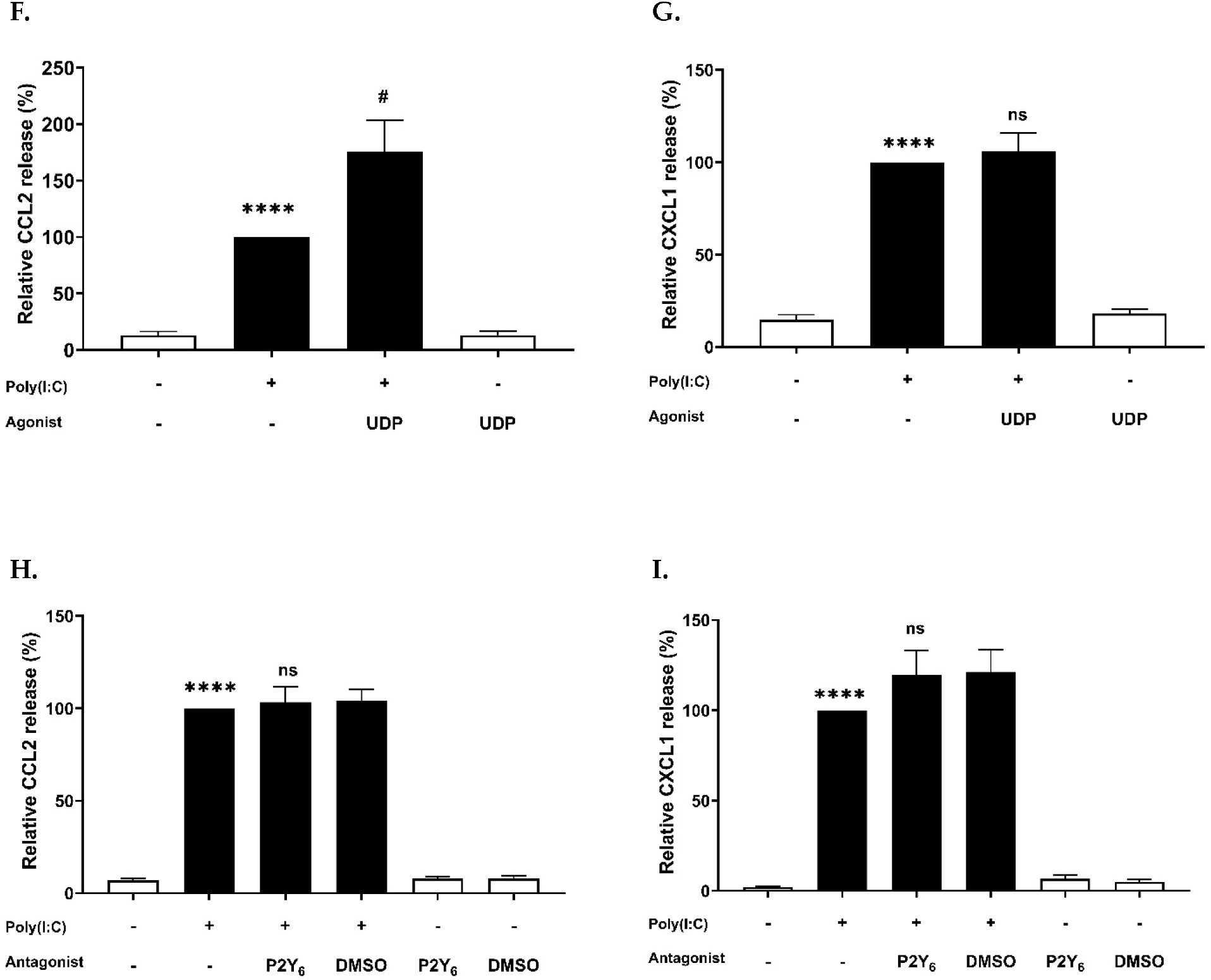
P2Y_6_ receptors are required for poly(I:C)-induced CCL2 and CXCL1 secretion in BMDMs. Basal expression of P2Y (Panel A) and P2X (Panel B) receptors in unstimulated BMDMs was determined by reanalysis of publicly available RNA-seq data from GSE198821 (PRJNA817065) and expressed as counts per million (CPM). Expression of *P2ry6, P2rx4*, and *P2rx7* was confirmed by RT-qPCR in BMDMs and normalized to *Gapdh* (Panel C). BMDMs were preincubated 20 minutes with the P2Y_6_ antagonist MRS2578 (5 µM), the P2X4 antagonist 5-BDBD (5 µM), the P2X7 antagonist A-438079 (5 µM), or the corresponding vehicle (DMSO 0.01%) for these three antagonists before stimulation with poly(I:C) (10 µg/mL). CCL2 (Panel D) and CXCL1 (Panel E) release into culture supernatants was measured by ELISA. Murine BMDMs were stimulated with a submaximal concentration of poly(I:C) (0.1 µg/mL) in the presence or absence of UDP (100 µM), and CCL2 (Panel F) and CXCL1 (Panel G) release was measured. BMDMs generated from *P2ry6*-deficient mice were stimulated with poly(I:C) (10 µg/mL) in the presence or absence of MRS2578 (5 µM) or the corresponding DMSO vehicle, and CCL2 (Panel H) and CXCL1 (Panel I) release were measured. All stimulations were performed for 18 h before ELISA analysis. For panels D–I, chemokine release is presented relative to poly(I:C) stimulation, which was set to 100 percent. The 100% values correspond to absolute CCL2 and CXCL1 concentrations of 1677 and 500 pg/mL for WT and 1428 and 356 pg/mL for *P2ry6*-deficient, respectively, following stimulation with poly(I:C) at 10 µg/mL (Panels D, E, H, and I), and 130 and 110 pg/mL, respectively, following stimulation with poly(I:C) at 0.1 µg/mL (Panels F and G). One symbol indicates *p* < 0.05, two symbols *p* < 0.01, and four symbols *p* < 0.0001. *, BMDMs stimulated with poly(I:C) versus unstimulated cells; #, BMDMs stimulated with poly(I:C) plus MRS2578 versus poly(I:C) plus DMSO (Panels D & E) or poly(I:C) plus UDP versus poly(I:C) alone (Panels F & G). For ns annotations, the comparison corresponds to poly(I:C) plus MRS2578 or UDP versus poly(I:C) alone.

To further assess whether P2Y_6_ activation modulates these responses, BMDMs were stimulated with a submaximal concentration of Poly(I:C) (0.1 µg/mL) in the presence of UDP (100 µM), the preferred endogenous agonist of P2Y_6_. UDP enhanced Poly(I:C)-induced CCL2 release (Fig. 4F), providing complementary evidence that P2Y_6_ positively supports CCL2 production. Previous studies have similarly reported that UDP/P2Y_6_ signaling can enhance inflammatory mediator production in macrophages, including chemokines (15). However, UDP did not significantly alter CXCL1 release, either alone or together with poly(I:C) (Fig. 4G).

Confirmation of MRS2578 target specificity was achieved using BMDMs from *P2ry6*-deficient mice. In these knockout cells, MRS2578 stimulation produced no significant change in poly(I:C)-induced CCL2 or CXCL1 release compared to the control (Fig. 4H, I). This lack of response demonstrates that the pharmacological actions of MRS2578 require functional P2Y_6_ expression. Collectively, these genetic and pharmacological data establish P2Y_6_ as a regulator of poly(I:C)-induced CCL2 and CXCL1 secretion, with opposing effects on these two chemokines.

### P2Y_6_ inhibition selectively modulates poly(I:C)-induced chemokine gene expression in BMDMs

Given the distinct effects of P2Y_6_ inhibition on CCL2 (Fig. 4D) and CXCL1 (Fig. 4E) secretion, we next examined whether these differences were also reflected at the transcriptional level. In addition to the expression of *Ccl2* and *Cxcl1*, we examined *Cxcl2*, another CXCR2 ligand involved in neutrophil recruitment (24,25), and *Cxcl10*, a distinct TLR3-responsive chemokine involved in the activation and recruitment of leukocytes such as T cells, NK cells and monocytes (26,27). *Cxcl10* was included to determine whether P2Y_6_ inhibition broadly amplified the TLR3 response or instead affected only selected chemokine programs.

Reanalysis of the RNA-seq dataset showed that, under basal conditions, *Cxcl10* was the most highly expressed chemokine examined, followed by *Ccl2* and *Cxcl2*, whereas *Cxcl1* expression was very low (Fig. 5A)(18). To determine whether the secretion phenotype was associated with transcriptional regulation, BMDMs were stimulated with poly(I:C) for 1 and 5 h in the presence or absence of MRS2578, and mRNA expression of *Ccl2, Cxcl1, Cxcl2*, and *Cxcl10* were measured by RT-qPCR (Fig. 5B-E).

**Figure 5.**
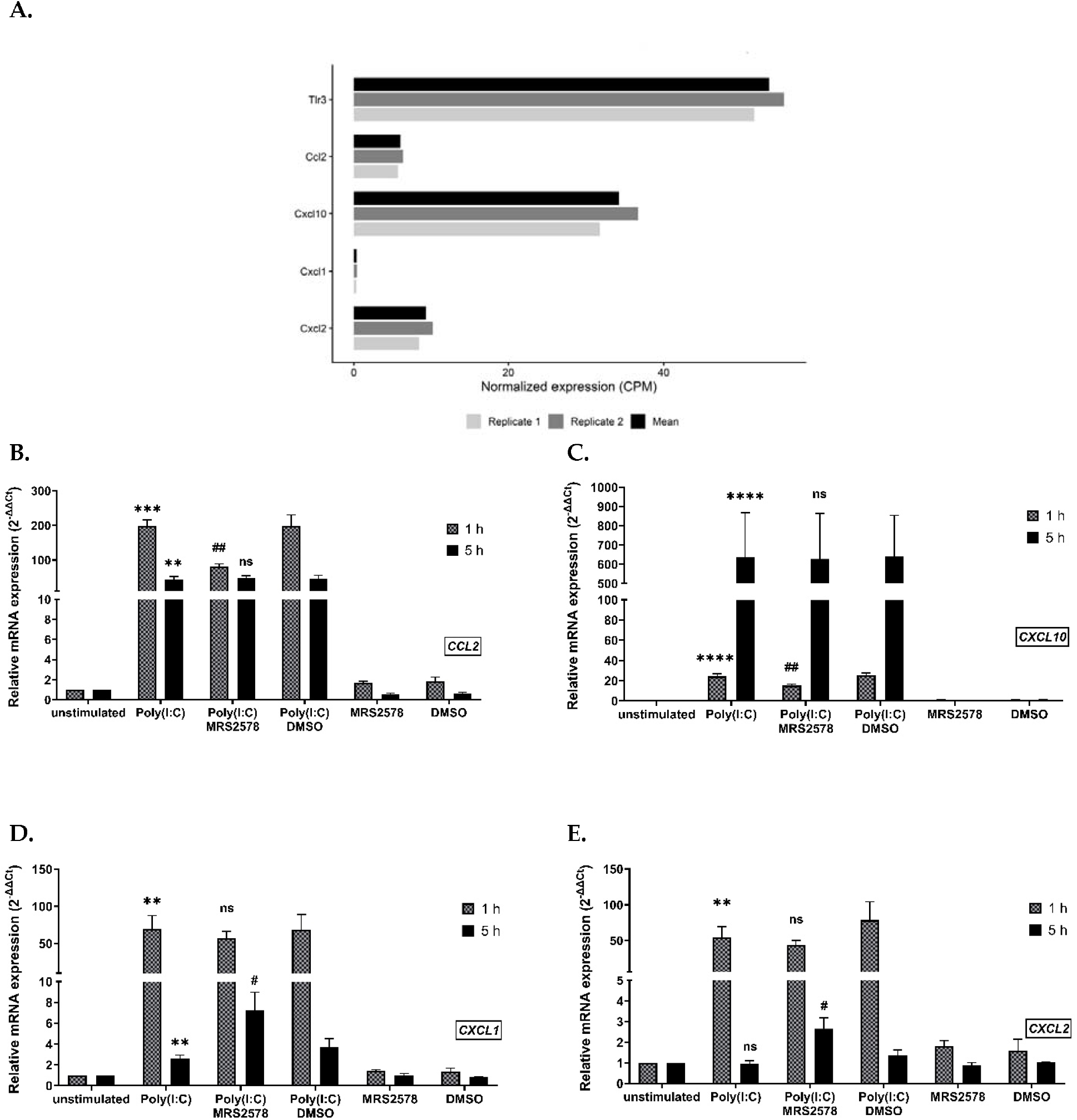
P2Y_6_ inhibition selectively modulates poly(I:C)-induced chemokine gene expression in BMDMs. Basal expressions of *Tlr3, Ccl2, Cxcl1, Cxcl2*, and *Cxcl10* in unstimulated BMDMs were determined by reanalysis of publicly available RNA-seq data from GSE198821 (PRJNA817065) and expressed as counts per million (CPM) (Panel A). BMDMs were pretreated with the P2Y_6_ antagonist MRS2578 (5 µM) or the corresponding DMSO vehicle (0.01%) before stimulation with poly(I:C) (1 µg/mL) for 1 (hatched grey bars) or 5 h (black bars) (Panels B-E). The expression of *Ccl2* (Panel B), *Cxcl10* (Panel C), *Cxcl1* (Panel D), and *Cxcl2* (Panel E) were assessed by RT-qPCR and normalized to *Gapdh*. Dat are presented as relative mRNA expression and represent the mean ± SEM of three independent experiments. One, two, three, and four symbols indicate *p* < 0.05, *p* < 0.01, *p* < 0.001, and *p* < 0.0001, respectively. *, BMDMs stimulated with poly(I:C) versus unstimulated cells; #, BMDMs stimulated with poly(I:C) plus MRS2578 versus poly(I:C) plus DMSO. For ns annotations, the comparison corresponds to poly(I:C) versus unstimulated cells when shown over the poly(I:C) bar, and poly(I:C) plus MRS2578 versus poly(I:C) plus DMSO when shown over the poly(I:C) plus MRS2578 bar.

Poly(I:C) induced distinct temporal patterns of chemokine expression, and the effects of P2Y_6_ inhibition were also time dependent. MRS2578 significantly reduced poly(I:C)-induced *Ccl2* and *Cxcl10* expression at 1 h, whereas these effects were no longer observed at 5 h (Fig. 5B and C). The early reduction in *Ccl2* expression is consistent with the decreased CCL2 secretion observed at 18 h (Fig. 4D).

In contrast, MRS2578 did not significantly alter poly(I:C)-induced *Cxcl1* or *Cxcl2* expression at 1 h but significantly increased both transcripts at 5 h (Fig. 5D and E). The later increase in *Cxcl1* expression is consistent with the enhanced CXCL1 secretion observed at 18 h (Fig. 4E). These findings indicate that P2Y_6_ signaling during poly(I:C) stimulation positively contributes to the early induction of *Ccl2* and *Cxcl10* while restraining the later *Cxcl1/Cxcl2* transcriptional response. Overall, P2Y_6_ inhibition differentially and temporally modifies poly(I:C)-driven chemokine responses rather than uniformly enhancing or suppressing inflammatory signaling.

## Discussion

In this study, we demonstrate that extracellular nucleotide signaling differentially regulates poly(I:C)-driven chemokine responses in murine BMDMs. Our findings further identify P2Y_6_ as an important component of this regulation, with distinct effects on CCL2 and CXCL1 responses. Overall, these results indicate that nucleotide signaling selectively shapes the chemokine profile generated following poly(I:C) stimulation.

Reanalysis of a publicly available RNA-seq dataset (Fig. 2A) (18), together with RT-qPCR validation in our murine BMDMs cultures (Fig. 2B), confirmed the expression of *Tlr3, Ccl2*, and *Cxcl1*. Moreover, poly(I:C) stimulation increased CCL2 and CXCL1 secretion in a dose-dependent manner (Fig. 2C and D), consistent with previous reports showing poly(I:C)-induced CCL2 and CXCL1 secretion in murine macrophages (28–30).

Having established this response, we next degraded extracellular nucleotides using apyrase. In the presence of poly(I:C), apyrase reduced CCL2 secretion while enhancing CXCL1 release (Fig. 3A-B), supporting a role for endogenous extracellular nucleotide signaling in the poly(I:C)-induced chemokine response. Consistently, suramin and RB2 also reduced poly(I:C)-induced CCL2 secretion (Fig. 3A). This pattern is consistent with evidence that extracellular ATP can differentially regulate chemokine expression in human keratinocytes, enhancing CCL2, CCL5, and CXCL8 while suppressing IFN-γ-induced CXCL9, CXCL10, and CXCL11 (31). Similarly, we previously demonstrated that apyrase reduced LPS-induced CCL2 secretion in human glioma cells (14). In our previous studies using human monocytes and monocytic cells, degradation of extracellular nucleotides with apyrase also reduced TLR4- and TLR1/2-induced CXCL8 secretion, indicating that endogenous nucleotide signaling can positively regulate this chemokine production in these cellular contexts (13,32).

To determine which P2 receptor contributed to this differential chemokine regulation, we next examined the nucleotide receptors most prominently expressed in murine BMDMs. Reanalysis of a publicly available RNA-seq dataset identified *P2ry6, P2rx4*, and *P2rx7* among the most highly expressed nucleotide receptor transcripts in murine BMDMs (Fig. 4A & B) (18). Expression that we have validated by RT-qPCR (Fig. 4C). This expression profile provided the rationale for investigating their contribution to poly(I:C)-induced chemokine responses. Among these receptors, only pharmacological inhibition of P2Y_6_ inhibition with the antagonist MRS2578 significantly altered poly(I:C)-induced chemokine secretion. MRS2578 reduced CCL2 while enhancing CXCL1 release, reproducing the pattern observed with apyrase (Fig. 4D and E). Moreover, MRS2578 produced no additional effect in *P2ry6*-deficient murine BMDMs (Fig. 4H and I), supporting this antagonist specificity on P2Y_6_ receptors. Our CCL2 findings are consistent with previous evidence from human glioma cells, where extracellular nucleotide depletion and P2Y_6_ blockade reduced LPS-induced CCL2 secretion (14). Similar contrasting effects on different chemokines have also been reported in rhesus macaque BMDMs, where MRS2578 enhanced TLR2/4-induced CXCL8 while reducing selected TLR5- and TLR8-induced cytokines (33). These findings indicate that P2Y_6_ effects are cells-, mediator- and PRR-dependent rather than uniformly proinflammatory.

The positive contribution of P2Y_6_ to CCL2 release was further supported by the ability of exogenous UDP to enhance CCL2 secretion in murine BMDMs stimulated with a submaximal concentration of poly(I:C)-(Fig. 4F). This agrees with previous evidence UDP–P2Y_6_ signaling enhance CCL2 expression, production and promote monocyte/macrophage recruitment during bacterial infection (34). More broadly, nucleotide-dependent CCL2 regulation is not restricted to P2Y_6_, as UTP acting predominantly through P2Y_2_ was shown to induce CCL2 expression and secretion in human THP-1 cells and macrophage-like cells (35). In contrast to its positive effect on CCL2, exogenous UDP did not alter CXCL1 secretion, despite the increased CXCL1 response observed following apyrase treatment and P2Y_6_ inhibition (Fig. 3B, 4E, and 5D) One possible explanation is that the P2Y_6_-dependent pathway restraining CXCL1 was already sufficiently engaged by endogenous nucleotides present during poly(I:C) stimulation, such that additional UDP produced little further effect. A similar pattern has been reported in human bronchial epithelial cells, where P2Y_6_ blockade altered stimulus-induced cytokine production, whereas addition of exogenous UDP to the inflammatory stimulus produced no further increase in IL-6 or IL-8. They interpreted this lack of additivity as evidence that the stimulus had already engaged an endogenous UDP–P2Y_6_ pathway (36). Likewise, in our study, the continued enhancement of CCL2 by exogenous UDP argues against global saturation of P2Y_6_ and suggests that P2Y_6_-dependent regulation differs among individual chemokine pathways. The effects of apyrase together with P2Y_6_ inhibition support the involvement of endogenous extracellular nucleotide release and signaling in this response.

To investigate the mechanisms underlying the differential regulation of these chemokines, we examined their transcriptional responses to P2Y_6_ inhibition following poly(I:C) stimulation. In addition to *Ccl2* and *Cxcl1*, we included *Cxcl2*, another CXCR2-binding neutrophil chemokine closely related to CXCL1 (24,25), and *Cxcl10*, a chemokine induced downstream of TLR3 that primarily mediates CXCR3-dependent recruitment of activated T cells, NK cells, and monocytes (26,27). Expression of these two latter chemokines in BMDMs was validated first by RNA-seq and subsequently validated by qPCR (Fig. 5A).

Poly(I:C) induced distinct kinetic profiles, with *Ccl2, Cxcl1*, and *Cxcl2* strongly induced at 1 h and declining by 5 h, whereas *Cxcl10* showed a delayed response and was markedly higher at 5 h (Fig. 5B–E). The rapid *Cxcl1/Cxcl2* response is consistent with early CXCL1 and CXCL2 transcription and de novo synthesis reported in TLR-stimulated macrophages (37) as well as early CXCL1 induction following poly(I:C) in vivo (38). Poly(I:C)-induced CXCL1 production has been shown to depend on TRIF, as CXCL1 release was abolished in TRIF-deficient murine cells following TLR3 stimulation (39). Others demonstrated in other cell types that inflammatory *Cxcl1/Cxcl2* expression is linked NF-κB- and STAT1-dependent pathways (40), although these mechanisms were not examined here. In contrast, the relatively sustained *Ccl2* expression observed at 5 h may reflect the autocrine IFN-β–IFNAR feedback mechanism described in poly(I:C)-stimulated BMDMs, which maintains *Ccl2* transcription after its initial induction (29). This secondary feedback may therefore explain why *Ccl2* declined less markedly than *Cxcl1* and *Cxcl2* between 1 and 5 h.

The effect of P2Y_6_ inhibition was similarly gene- and time-dependent. MRS2578 reduced poly(I:C)-induced *Ccl2* and *Cxcl10* expression at 1 h but not at 5 h (Fig. 5B and C), suggesting that P2Y_6_ contributes mainly to their early induction. The early decrease in *Ccl2* may therefore contribute to the reduced CCL2 secretion observed at 18 h. Additional post-transcriptional regulation of CCL2 also remains possible, as CCL2 production can be controlled (41) through mRNA-associated mechanisms and intracellular trafficking/secretion pathways (42). Consistent with this finding, our previous study showed that acute P2Y_6_ inhibition reduced poly(I:C)-induced CXCL10 expression and secretion in murine intestinal epithelial cells (17). In contrast, MRS2578 did not alter the early *Cxcl1/Cxcl2* response but resulted in higher transcript levels at 5 h, when expression induced by poly(I:C) alone had markedly declined (Fig. 5D and E). This pattern suggests that P2Y_6_ may limit the persistence of the *Cxcl1/Cxcl2* mRNA response rather than their initial induction. The higher *Cxcl1* transcript level at 5 h may reflect prolonged transcription and/or increased mRNA stability and may contribute to the enhanced CXCL1 accumulation detected at 18 h. Thus, P2Y_6_ appears to support early *Ccl2/Cxcl10* responses while restraining the later *Cxcl1/Cxcl2* program, although the downstream mechanisms remain to be defined.

The opposing effects of P2Y_6_ on CCL2 versus the CXCL1/CXCL2 program may influence both the composition and recruitment of leukocyte during TLR3-driven inflammation (22,43). CCL2–CCR2 signaling promotes inflammatory monocyte recruitment and macrophage accumulation in atherosclerosis (44) and experimental autoimmune encephalomyelitis (45). Elevated CCL2 has been reported in rheumatoid arthritis (46). In contrast, CXCL1 and CXCL2 signals through CXCR2 promote neutrophil recruitment (5), which can contribute to tissue injury during acute lung inflammation (47) and influenza pneumonitis (48). CXCL10 represents an additional CXCR3-associated inflammatory program, indicating that P2Y_6_ regulation extends beyond a simple monocyte-versus-neutrophil response. Whether these chemokine changes translate into altered leukocyte recruitment would be interesting to examine in appropriate *in vivo* models.

This context dependence is not unique to P2Y_6_, as similar complexities have been observed when targeting other inflammatory pathways. TNF neutralization unexpectedly exacerbated multiple sclerosis, with increased inflammatory activity and worsening clinical disease in patients receiving anti-TNF treatment (49,50). Similarly, blockade of IL-17 receptor signaling with brodalumab was associated with worsening Crohn’s disease and led to early termination of the clinical trial (51). Experimental studies further showed that inhibition of IL-17A/IL-17RA can aggravate intestinal inflammation by impairing epithelial barrier integrity (52). These examples illustrate that inflammatory pathways can exert both pathogenic and protective functions depending on the biological context. Accordingly, defining the mechanisms through which P2Y_6_ selectively regulates individual chemokine pathways may provide a more precise therapeutic approach than global inhibition of the receptor.

## Conclusion

Our data reveals a previously unrecognized role for nucleotide signaling in shaping the chemokine profile generated during TLR3 activation in macrophages. The novelty of this study lies not only in demonstrating the involvement of P2Y_6_ in a poly(I:C)-driven PRR response, but also in showing that the same receptor may exert opposing effects on distinct chemokine programs downstream of the same stimulus. P2Y_6_ may indeed regulate inflammatory responses in a gene- and time-dependent manner rather than functioning as a uniformly pro- or anti-inflammatory receptor. This selective regulation may influence the composition of the inflammatory response and highlights the potential value of targeting specific downstream pathways rather than globally inhibiting P2Y_6_.

## Author Contributions

**Fatemeh Salarpour:** Conceptualization; Investigation; Formal analysis; Writing—original draft, review and editing. **Anaïs Brafine:** Investigation (preliminary experiments); Formal analysis. **Ashley Charre:** Investigation (preliminary experiments); Formal analysis. **Julie Pelletier:** Resources (Materials); Writing—review and editing. **Jean Sévigny:** Conceptualization; Supervision; Writing—review and editing.

## Institutional Review Board Statement

Not applicable. Informed

## Consent Statement

Not applicable.

## Data Availability Statement

The original contributions presented in this study are included in the article. Further inquiries can be directed at the corresponding author(s).

